# Extravascular spaces are the primary reservoir of antigenic diversity in *Trypanosoma brucei* infection

**DOI:** 10.1101/2022.06.27.497797

**Authors:** Alexander K. Beaver, Zhibek Keneskhanova, Raúl O. Cosentino, Brian L. Weiss, Erick O. Awuoche, Gretchen M. Smallenberger, Gracyn Y. Buenconsejo, Nathan P. Crilly, Jaclyn E. Smith, Jill M.C. Hakim, Bailin Zhang, Bryce Bobb, Filipa Rijo-Ferreira, Luisa M. Figueiredo, Serap Aksoy, T. Nicolai Siegel, Monica R. Mugnier

## Abstract

The protozoan parasite *Trypanosoma brucei* evades clearance by the host immune system through antigenic variation of its dense variant surface glycoprotein (VSG) coat, periodically “switching” expression of the VSG using a large genomic repertoire of VSG-encoding genes^1–6^. Recent studies of antigenic variation in vivo have focused near exclusively on parasites in the bloodstream^4,7,8^, but research has shown that many, if not most, parasites reside in the interstitial spaces of tissues^9–13^. We sought to explore the dynamics of antigenic variation in extravascular parasite populations using VSG-seq^7^, a high-throughput sequencing approach for profiling VSGs expressed in populations of *T. brucei*. Here we show that tissues, not the blood, are the primary reservoir of antigenic diversity during both needle- and tsetse bite-initiated *T. brucei* infections, with more than 75% of VSGs found exclusively within extravascular spaces. We found that this increased diversity is correlated with slower parasite clearance in tissue spaces. Together, these data support a model in which the slower immune response in extravascular spaces provides more time to generate the antigenic diversity needed to maintain a chronic infection. Our findings reveal the important role that extravascular spaces can play in pathogen diversification.

## Main

Every pathogen must contend with the adaptive immune response of its host. *Trypanosoma brucei,* a protozoan parasite and causative agent of human and animal African Trypanosomiasis, has evolved a sophisticated strategy to evade this highly flexible and specific host response^14^. Transmitted by the bite of the tsetse fly, *T. brucei* lives extracellularly in the blood, lymph, and interstitial tissue spaces of its mammalian host^15^. To escape clearance by a continuous onslaught of host antibodies, the parasite periodically “switches” expression of its immunogenic variant surface glycoprotein (VSG) coat to new, antigenically distinct variants^1^. With a genomic repertoire of thousands of different VSG-encoding genes^2,3,5,6^ and the ability to generate novel VSGs through recombination^4,8,16^, the parasite has an enormous capacity for altering its antigenic profile.

Studies examining *T. brucei* antigenic variation in vivo have focused nearly exclusively on parasites in the blood, revealing complex VSG expression dynamics^4,7,8^. However, it has recently become clear that many, if not most, *T. brucei* parasites inhabit extravascular spaces during both experimental and natural infections^9–13^. Though research has shown that tissue-resident parasites adapt to these environments^11^, cause tissue-specific symptoms^15^, and are associated with increased disease severity^13^, it remains unclear why parasites invade tissue spaces and what role these populations might play in infection.

Several older studies suggested a role for tissue-resident parasites in antigenic variation. These studies found that brain- and lymph-resident parasite populations were antigenically distinct from those in the blood^17–20^, and that antigenic types detectable in the lymphatic fluid could be detected in the blood at later timepoints during infection^19^. This led these researchers to hypothesize that extravascular spaces might be a site for antigenic variation, with antigenic variants generated in extravascular spaces contributing to systemic immune evasion. However, a later study focusing on very early timepoints post-infection found no antigenic difference between *T. brucei* populations within the blood and several extravascular spaces^21^, and the community consequently ruled out a role for extravascular parasites in antigenic variation^22,23^. These early investigations were limited by the methodology available at the time, which relied on the use of VSG-specific antisera to analyze parasite antigenic diversity. Modern high-throughput sequencing methods allow VSG expression to be measured accurately and in high resolution^7^. Given the mounting evidence that *T. brucei* parasites persist in and adapt to extravascular spaces, there is a clear need to reinvestigate the dynamics of VSG expression within tissue spaces.

Here, we use VSG-seq^7^, a targeted RNA-sequencing approach for profiling the VSGs expressed in *T. brucei* populations, to characterize the VSGs expressed by extravascular *T. brucei* parasites. Our results show that extravascular spaces are major reservoirs of antigenic diversity during *T. brucei* infection and that this parasite niche is central to the parasite’s ability to continuously outmaneuver the immune system.

### Extravascular spaces contain most of the VSG diversity

To investigate how tissue-resident parasites contribute to antigenic variation in vivo, we intravenously (IV) infected 12 mice, each with ∼5 pleomorphic *T. brucei* EATRO1125 90-13 parasites^24^. We collected blood, then perfused mice with PBS-glucose (Extended Data Fig. 1) and harvested the heart, lungs, gonadal fat, subcutaneous fat, brain, and skin at 6, 10, and 14 days post-infection. For each sample, we extracted RNA and quantified *T. brucei* VSG expression using VSG-seq^7^. A single “initiating” VSG (either AnTat1.1 or EATRO1125 VSG-421) dominated expression in both the blood and tissues on day 6 (Fig. 1a), in line with previous observations^21,25^. At later time points, VSG expression dynamics became more complex, with more VSGs expressed, a unique composition of VSGs in each tissue, and no single dominating variant (Fig. 1a). Although tissue-specific expression of variant surface proteins is a feature in other organisms that use antigenic variation^26–31^, we found no evidence for tissue-specific VSGs or VSG sequence motifs (Extended Data Fig. 2). Instead, we observed an increase in antigenic diversity in extravascular spaces. The number of detectable VSGs in tissue spaces was, on average, two to four times higher than the blood (Fig. 1c). This did not correlate with parasite load (Extended Data Fig. 3 and Extended Data Fig. 4ab) and was not driven by any specific tissue (Fig. 1d). In addition, parasite differentiation to the non-dividing tsetse infective form via quorum sensing, which is marked by expression of the PAD1 gene^32^, did not correlate with VSG diversity in either the blood or tissues at the population level (Extended Data Fig. 4c). The overall contribution of tissue-resident parasites to antigenic diversity in a single infection was large: at any time, ∼87% of expressed VSGs in any individual infection were found exclusively within extravascular spaces (Fig. 1b).

**Fig. 1.**
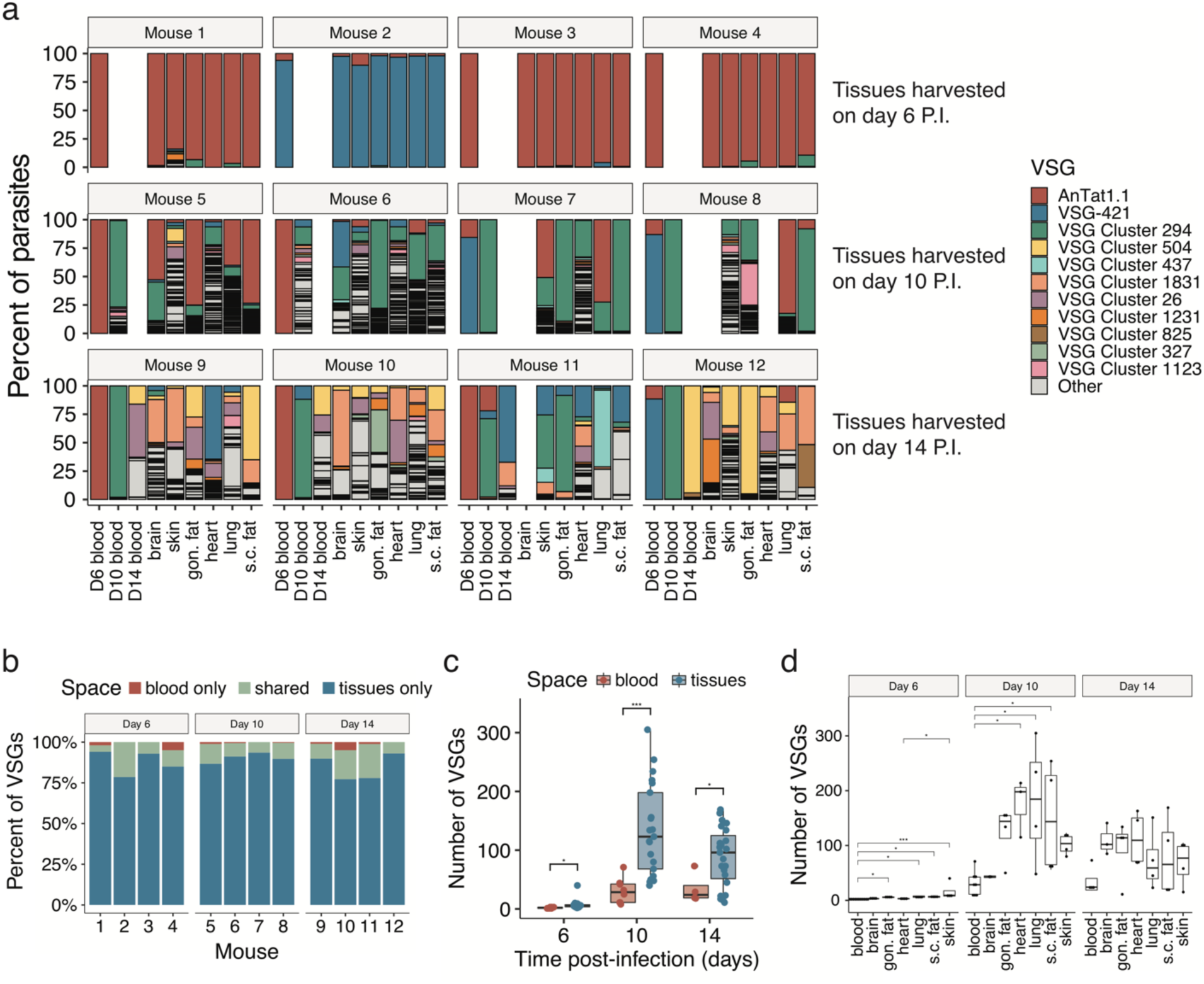
Extravascular parasites harbor most of the antigenic diversity in an infection. (**a**) The percentage of parasites expressing each VSG within a space. The 11 VSGs with the highest overall expression are colored, and all other VSGs are in grey as “other”. Gon. fat = gonadal fat, s.c. fat = subcutaneous fat, P.I. = post-infection. (**b**) Stacked bar graphs from each infected mouse representing the percentage of VSGs that were found exclusively within the blood (red), exclusively within tissue spaces (blue), or shared by both the blood and at least one tissue (green). (**c**) Quantification of the number of VSGs found within the blood (red) or tissue spaces (blue) at each time point (Shapiro-Wilk normality test followed by a two-tailed Student’s t-test BH corrected). (**d**) The number of VSGs in each tissue space (Shapiro-Wilk normality test followed by a two-tailed Dunnett’s test) (For statistical tests *P<0.05, ***P<0.001).

VSG-seq is a bulk measure of VSG expression. To be sure that the increased VSG diversity we observed in tissues was not due to the derepression of silent VSGs within individual cells, we performed single-cell RNA sequencing using the SL-Smart-seq3xpress platform to quantify VSG expression in single cells from blood and tissue samples^33^. Because *T. brucei* can form new “mosaic” VSGs through recombination^4,8,16,34^, which are, by definition, absent from reference genomes, we initially used our VSG-Seq pipeline to *de novo* assemble VSGs in each cell. This approach accounts for the possibility that expressed VSGs might be absent from the reference genome, either as a result of mosaic VSG formation or as a result of an incomplete genome assembly, potentially affecting quantification and/or read mapping. Only one VSG assembled in most cells (94.2%; Extended Data Fig. 5a). We also mapped sequencing reads to the EATRO1125 genome, which could reveal more subtle signatures of derepression. By this analysis, there was no obvious difference between the blood and the tissues in the number of expressed VSGs in each cell (Extended Data Fig. 5b) or in the relative expression of the most abundant VSG in each cell (Extended Data Fig. 5c).

To estimate the number of cells maintaining monogenic expression, we defined a cell as maintaining monogenic expression if 80% of UMIs mapping to VSGs mapped to a single VSG. Although mapping to the genome suggested that most cells (62.7%) maintained monogenic expression based on this threshold, the proportion of cells estimated to maintain monogenic expression was lower than estimated by *de novo* assembly. Comparison of the two analyses revealed that of those cells expressing >1 VSG by genome alignment, 85% were found to express only one VSG by mapping to the *de novo* assembled VSGs and using the same 80% threshold for defining monogenic expression. Further investigation revealed that in most of these cases (95.3%) the alignment to multiple genomic VSGs was an artifact, where the assembled VSG was not well represented within the annotated sequences of the EATRO1125 genome or there were several VSGs with high similarity to the assembled VSG, leading to inaccurate VSG expression quantification (Extended Data Fig. 5d). We estimate that 93.4% of cells were likely expressing only one VSG, with no bias for multigenic VSG expression in tissue spaces (Extended Data Fig. 5e, shades of green and Extended Data Table 2). In 5.2% of cells, reads mapped to multiple EATRO1125 VSGs, but no VSG could be assembled (grey). While it is impossible to distinguish between multi- and monogenic VSG expression in this set of cells, the proportion of cells in this category did not differ between blood and tissues. The few cells that appear to express multiple VSGs (1.05% of cells in the blood and 1.03% of cells in the tissues) could indicate sorting doublets or, more interestingly, could represent cells mid-switch (Extended Data Fig. 5e, red and purple). Overall, these data suggest that VSG monogenic expression is maintained by most cells in both extravascular spaces and the blood and that the increased VSG diversity we observe in tissue spaces is unlikely to be due to the specific derepression of silent VSGs in tissue populations.

### Frequently expressed VSGs first appear in tissue populations

The high antigenic diversity observed in tissues could serve to maintain a chronic infection. If antigenic variation occurs relatively rarely in the bloodstream, then parasites from extravascular spaces might serve as a source of new, antigenically distinct, VSGs. In line with this, tissue spaces contain more “unique VSGs”, those VSGs that are expressed exclusively within one space in an infection, than the blood (Fig. 2ab). To examine the potential for tissue-resident VSGs to contribute to antigenic variation systemically, we identified VSGs only expressed in tissues on day 6 post-infection and analyzed whether they later appeared within the blood. The majority (74%) of these VSGs were expressed within the blood on day 10 or 14. Analysis of individual VSGs revealed that rare VSGs expressed at low levels exclusively in tissue spaces also have the potential to become ubiquitously expressed within a host (Fig. 2c). In addition to a model in which tissue spaces provide new VSGs to re-seed the blood, it is possible that tissue-resident parasites undergo antigenic variation before blood-resident populations. Therefore, these data could be explained by either a trigger within the tissue environment that induces parasite switching or a differential selective pressure imposed by the tissue environment.

**Fig. 2.**
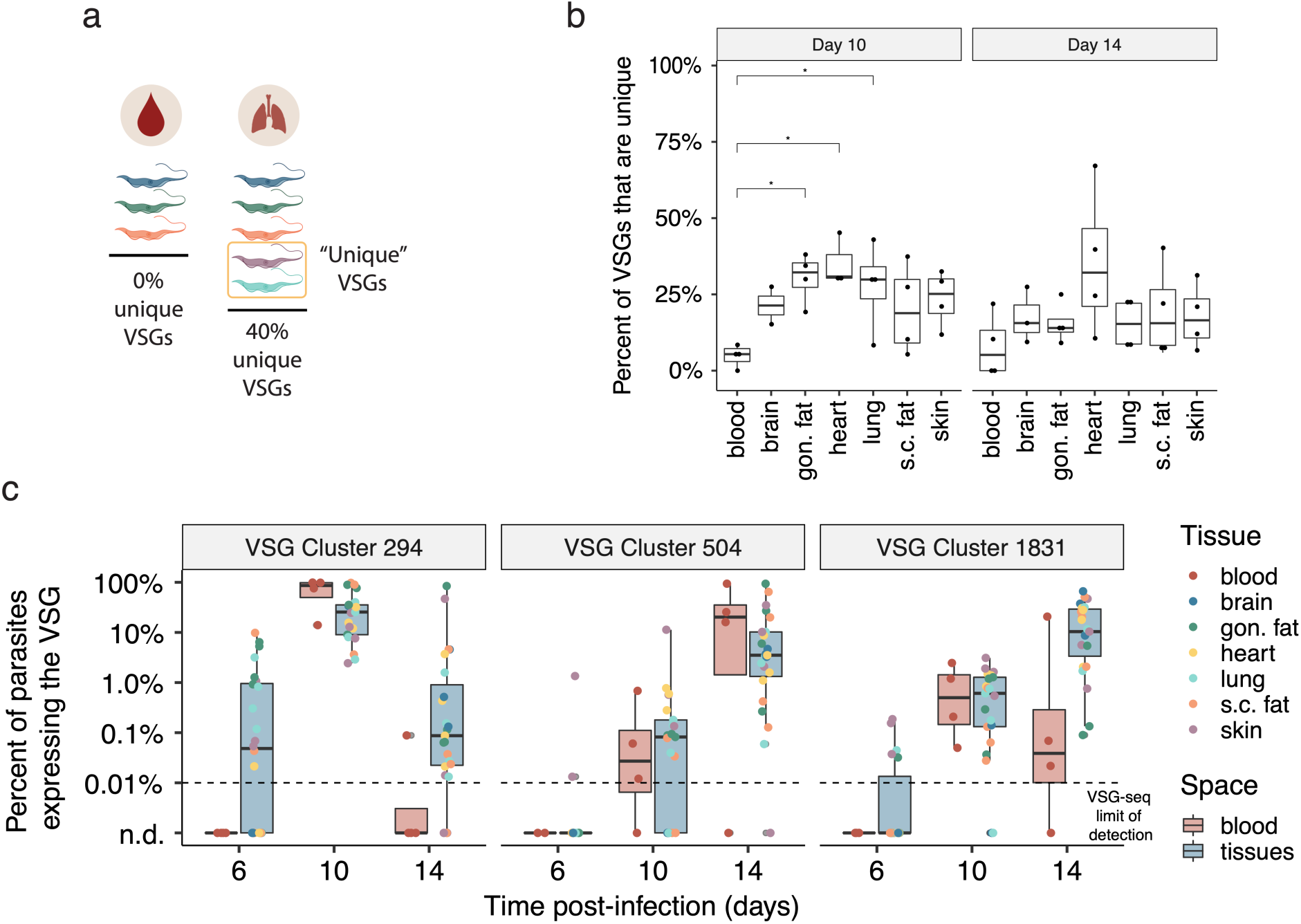
Tissue-resident parasites express a unique repertoire of VSGs during infection. (**a**) We define “unique” VSGs as those VSGs solely found within a specific space in a mouse (Created with BioRender.com). (**b**) The percentage of VSGs that were unique to one space within a mouse (Shapiro-Wilk normality test followed by a two-tailed Dunnett’s test, *P<0.05). Day 6 samples were excluded from this analysis because few VSGs are expressed at this point. Gon. fat = gonadal fat, s.c. fat = subcutaneous fat. (**c**) The expression of three representative VSGs (cluster 294, 504, and 1831) within blood and tissue samples on days 6, 10, and 14. “n.d” indicates that the VSG was not detected.

### VSG-specific parasite clearance is delayed in tissues compared to the blood

Indeed, our data suggest that the environment within tissue spaces is distinct from the blood, with parasite clearance occurring at different rates in each space. While parasites expressing the initiating VSG were cleared from the blood by day 10 post-infection, they were not cleared from tissues until at least day 14 (Fig. 3a). This suggests that VSG-specific parasite clearance from extravascular spaces is delayed, but not abolished, compared to the blood. VSG-seq is a measure of VSG expression at the transcript level, however. To confirm this observation at the protein level, we performed flow cytometry on *T. brucei* cells from the blood, lungs, and gonadal fat, using the tdTomato-expressing “triple reporter” *T. brucei* EATRO1125 AnTat1.1E cell line^35^ (Fig. 3bc). The flow cytometry analysis showed a detectable AnTat1.1-expressing tissue parasite population at day 13 post-infection, a timepoint at which AnTat1.1^+^ parasites were undetectable, or nearly undetectable, in the blood. Similar to the increase in antigenic diversity we observed in every tissue space, this delay in clearance, observed at both the RNA and protein levels, was not tissue-specific. Thus, the immune mechanisms influencing extravascular parasite clearance appear to be general features of extravascular spaces.

**Fig. 3.**
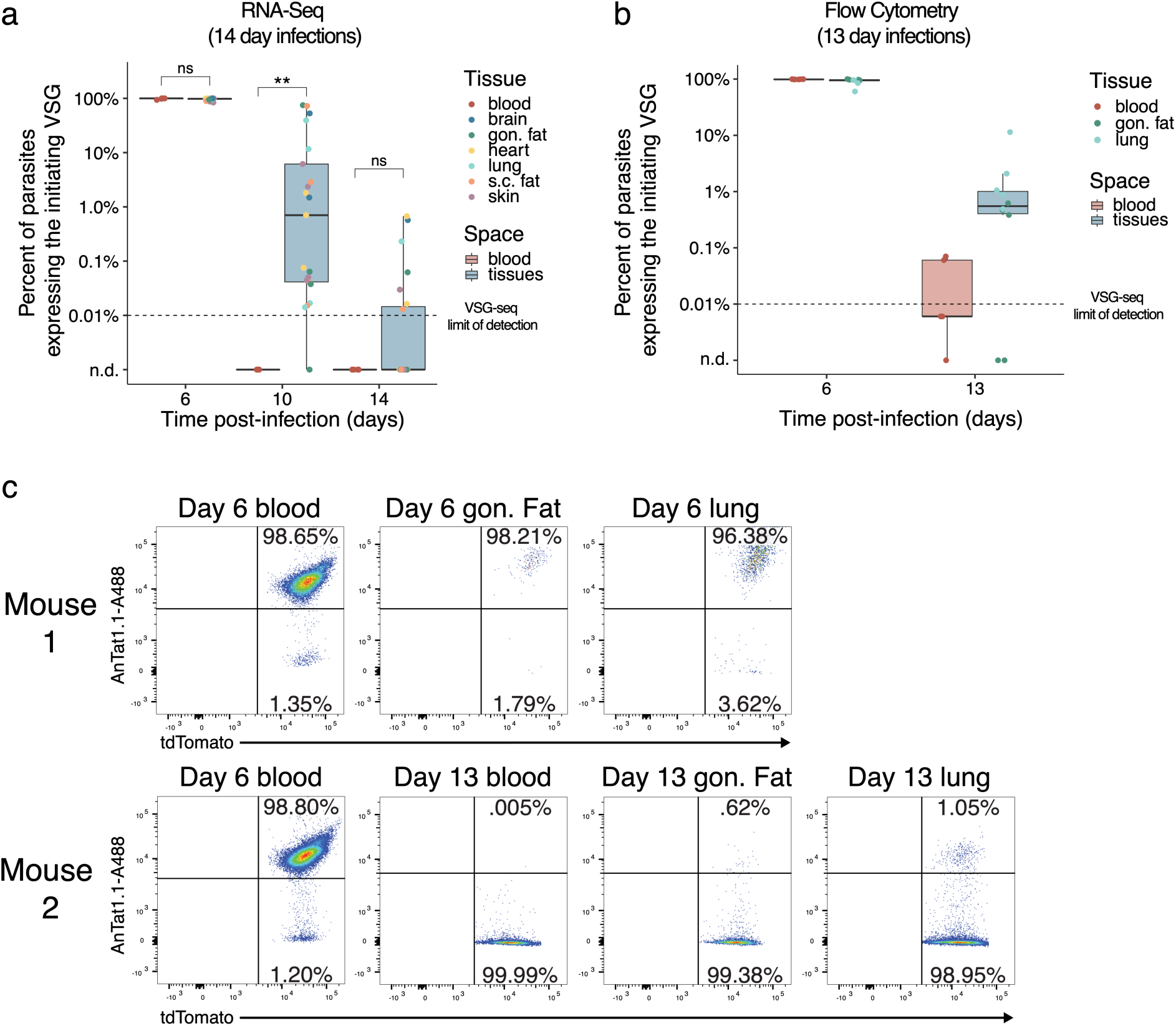
VSG-specific parasite clearance is slower in tissues than in the blood. (**a**) The percentage of parasites expressing the initiating VSG (AnTat1.1 or VSG-421) at days 6, 10, and 14 post-infection. Tissue samples were grouped together (blue) and compared to blood samples (red) (two-tailed Wilcoxon test, ns = not significant, **P<0.01). “n.d” = not detected, gon. fat = gonadal fat, s.c. fat = subcutaneous fat. (**b**) Quantification of the number of parasites that were tdTomato positive and stained positive for AnTat1.1 by flow cytometry (n = 5 mice). The horizontal dotted line represents the limit of detection for VSG-seq. “n.d” indicates that the VSG was not detected. (**c**) Representative flow cytometry plots from tissues collected from mice infected with chimeric triple marker parasites that express tdTomato constitutively in their cytoplasm. Parasites were stained with anti-AnTat1.1 antibody.

### Infections initiated by tsetse fly bite also show increased diversity and delayed clearance in extravascular spaces

A benefit of starting infections with a small intravenous inoculum is that it creates convenient and reproducible infections. In nature, however, a tsetse fly bite introduces thousands of parasites into the skin, each expressing a single metacyclic VSG (mVSG) from a limited repertoire^36^. To ensure that our observations held true in this more complex context, we repeated our infections using a more natural tsetse bite infection model. Infections were initiated in 5 mice by tsetse bite using flies infected with RUMP 503 *T. brucei* parasites^37^. We used VSG-seq to quantify VSG expression in the blood on day 5 post-infection and the blood and tissues on day 14 post-infection. In line with our previous observations, we found that in tsetse-initiated infections ∼80% of VSGs were exclusively expressed within extravascular spaces (Fig. 4a) and tissue populations harbored more VSGs than the blood (Fig. 4b). This demonstrates that extravascular spaces are the primary reservoir of antigenic diversity, even when infections are initiated by fly bite.

**Fig. 4.**
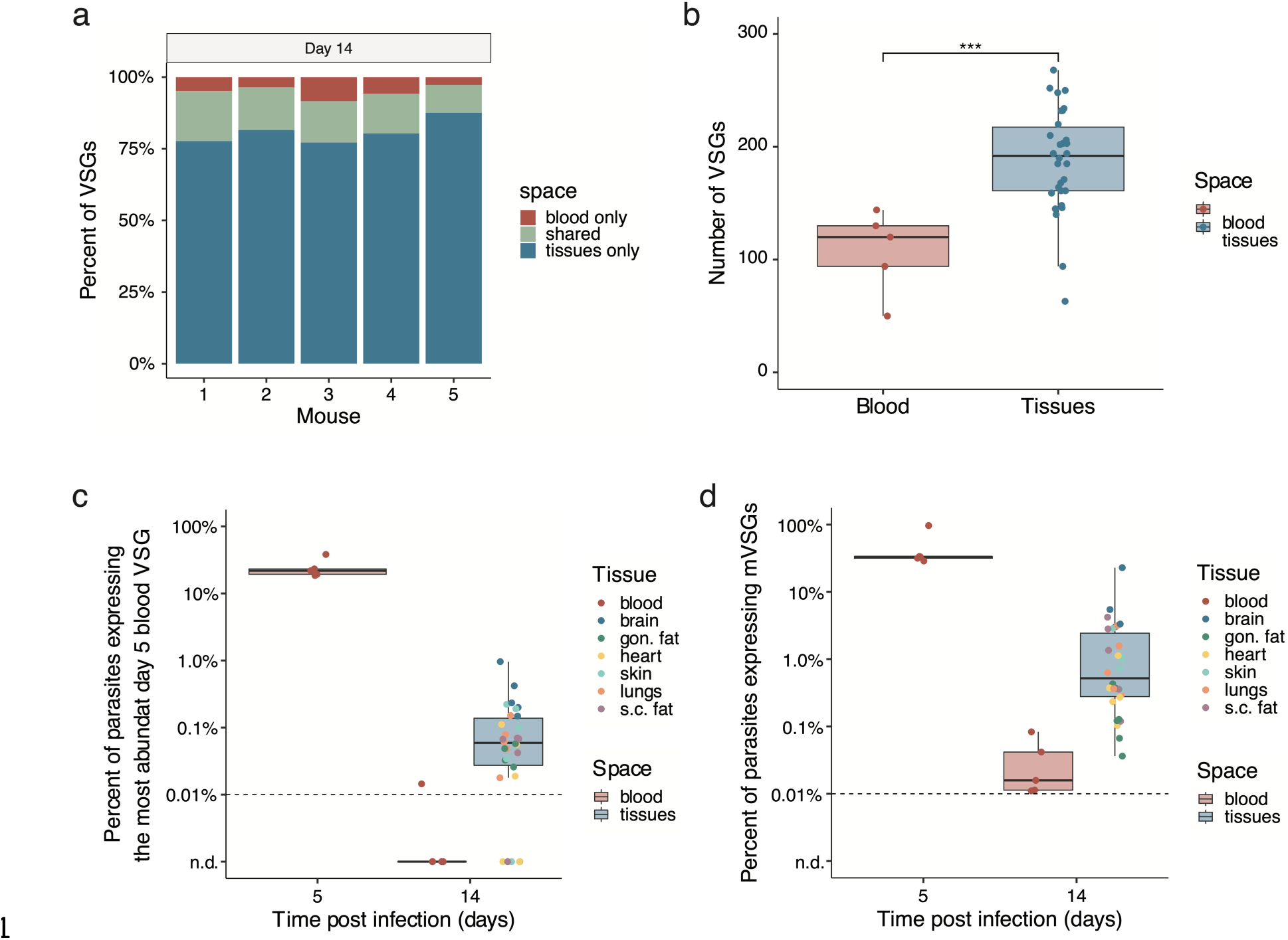
Tsetse bite-initiated infections show increased antigenic diversity and delayed immune clearance in extravascular spaces. Data from five mice infected with RUMP 503 parasites from a tsetse fly bite. (**a**) Bar graphs representing the percentage of VSGs in each mouse that were found exclusively within the blood (red), exclusively within tissue spaces (blue), or shared by both the blood and at least one tissue (green). (**b**) Quantification of the number of VSGs found within the blood (red) or tissue spaces (blue) on day 14 post-infection (Shapiro-Wilk normality test followed by a two-tailed Student’s t-test BH corrected, ***P<0.001). (**c**) The percentage of parasites within each mouse expressing the most abundant VSG from the day 5 blood. (**d**) The percentage of parasites expressing one of the mVSGs found to be expressed by RUMP 503 parasites in the salivary gland of a tsetse fly.

Parasite populations were more diverse at early time points in tsetse infections than IV infections, likely due to the larger and more heterogeneous inoculum delivered by the fly. To analyze VSG-specific parasite clearance in tissues, we quantified expression of the most abundantly expressed VSG in the blood of each mouse on day 5 as well as the known mVSG repertoire from this parasite strain, which should represent the repertoire of VSGs expressed at the start of an infection. In both cases, we found that on day 14, tissue spaces still contained parasites expressing the most abundant VSG and/or mVSGs, while these VSGs were expressed by no or very few parasites in the blood (Fig. 4c,d). This suggests that in tsetse-bite-initiated infections, as we observed in IV infections, tissue parasite populations are cleared at a slower rate than parasites in the blood.

### Delayed clearance of parasites leads to increased VSG diversity

The increased antigenic diversity in extravascular spaces could be explained by the distinct clearance dynamics we observe, as prolonged survival in tissues could provide more time for parasites to switch. To test whether there was a link between parasite survival and increased diversity, we sought to interrupt parasite clearance in tissues. Because parasite clearance in the blood coincides with the appearance of anti-VSG IgM, between days 8 and 10, and parasite clearance in the tissues coincides with the anti-VSG IgG response, between days 10 and 14^1,12,38–41^, we hypothesized that clearance in tissues is dependent on the anti-VSG IgG response. Thus, the loss of IgG might abrogate the clearance of tissue-resident parasites. To test this hypothesis, we infected activation-induced cytidine deaminase (AID) knockout (AID Cre) mice^42^, which only produce IgM antibodies (Extended Data Fig. 6), and analyzed blood and tissues from days 6 and 14 post-infection by VSG-seq. As expected, clearance of the initiating VSG was severely delayed in tissues on day 14 in AID^-/-^ mice, suggesting that IgG is important, if not critical, for the clearance of extravascular parasites. This could be explained by the fact that IgM, a bulky pentamer, does not diffuse efficiently into tissue spaces^43,44^, while IgG, a monomer, readily diffuses. We also observed a defect in the clearance of blood-resident parasites in AID^-/-^ mice compared to the blood of WT mice (Fig. 5a). In both the blood and tissues of AID^-/-^ mice, where clearance was delayed, more VSGs were detected on day 14 post-infection compared to wild-type (Fig. 5b), revealing a direct relationship between the timing of parasite clearance and VSG diversity. Regardless of their local environment (intra- or extravascular), longer-lived parasite populations generated more diverse sets of VSGs.

**Fig. 5.**
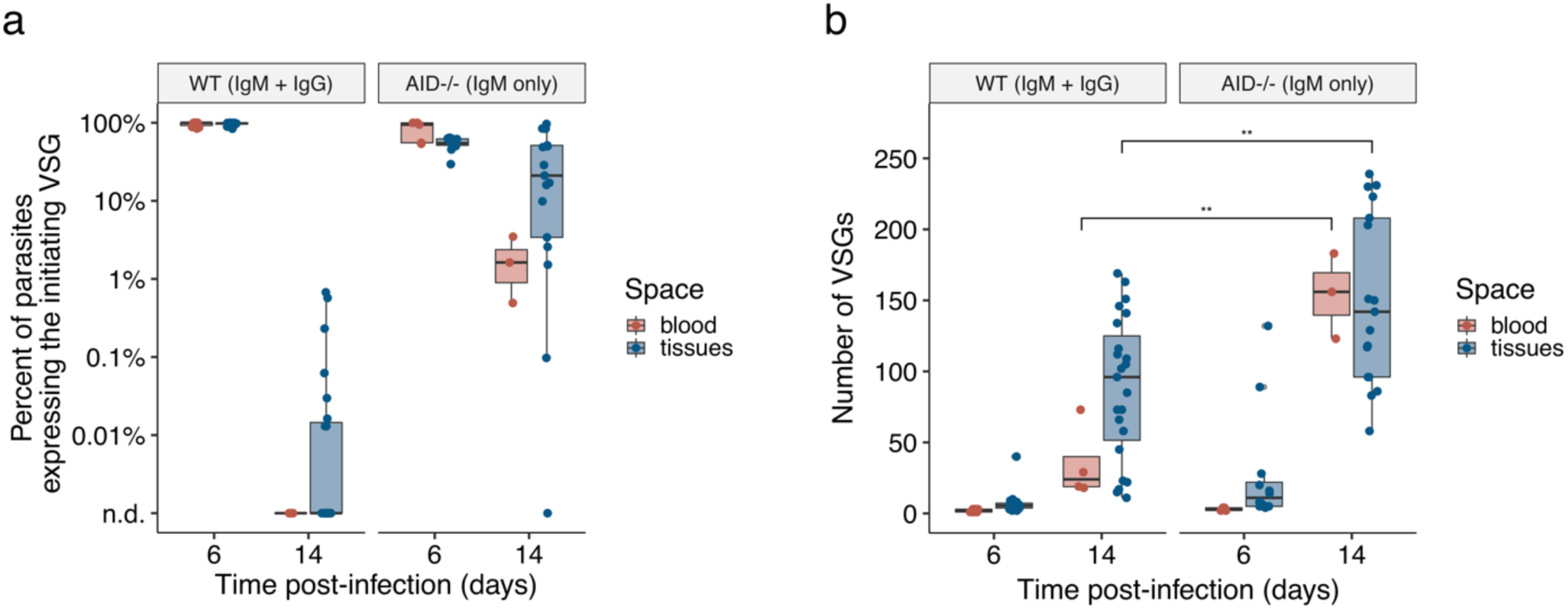
Delayed parasite clearance correlates with an increase in VSG diversity. (**a**) The percentage of parasites expressing the initiating VSG (AnTat1.1 or VSG-421) in both WT and AID^-/-^ mice. “n.d” indicates that the VSG was not detected. (**b**) The number of VSGs expressed within the blood (red) and tissues (blue) of WT and AID^-/-^ mice (Shapiro-Wilk normality test followed by a two-tailed Student’s t-test, **P<0.01).

## Discussion

The idea that antigenic variation might occur outside of the bloodstream, with extravascular populations contributing to immune evasion within the bloodstream, is not a new one. Here, using modern high-resolution techniques, we provide evidence for this long-standing hypothesis. In both needle- and tsetse bite-initiated infections, we find that extravascular spaces are the primary reservoir of VSGs, accounting for the vast majority of antigenic diversity in any individual infection. The number of VSGs we detected in the blood matches previous estimates^4,7,8^, while the diversity in tissues is two to four times higher. Our data suggest that this is at least partially due to slower clearance dynamics in extravascular spaces and highlight the role that tissue spaces can play in pathogen diversification.

Although extravascular parasite populations were highly antigenically diverse, we saw no evidence of tissue-specific VSG expression. Because parasites invade tissues efficiently before much VSG switching has occurred, it appears unlikely that any specific VSG is required for tissue invasion. Whether VSGs could influence parasite fitness in specific host spaces is less clear. We measured VSG expression up to day 14 post-infection, at which point tissue-resident populations are just beginning to diverge from one another and the blood. It is therefore possible that, as these populations further evolve, there may be a selection for VSGs better adapted to certain tissue spaces.

It has previously been shown that in the bloodstream alone *T. brucei* expresses more VSGs than appear to be required for immune evasion^7^. Here we find that the expressed VSG diversity within host tissues is even greater. While on the surface it could appear to be disadvantageous for *T. brucei* to use so many different antigens this quickly, this striking diversity could be important for the parasite. In natural infections, particularly in wild animals where pre-existing anti-VSG immunity is more likely to exist, a high switch rate may be required to ensure some parasites successfully evade the host’s existing antibody repertoire. Moreover, infections in the wild can last for months to years^14,45^. During these long infections, the large reservoirs of VSGs found in tissues may be essential for the maintenance of a chronic infection.

Indeed, our data support the idea that the large reservoirs of antigenic diversity in extravascular spaces contribute to systemic infection when parasites re-enter the blood after switching: rare VSGs expressed exclusively in tissues at early time points are expressed in the blood and other spaces later. This is also in line with another recent study which showed that blood-resident parasites are largely non-replicative, indicating that tissue-resident parasites may be required to re-seed the blood^46^. There is another intriguing explanation for this observation, however. Switching in *T. brucei* is known to be semi-predictable^7,47^, and it is possible that tissue-resident parasites simply switch earlier, or more frequently, than those in the blood. In this case, the same VSGs would arise independently in every population without any parasite movement between spaces. While the high vascular permeability observed after the initial stages of infection indicates that parasites likely move back and forth between the vasculature and extravascular spaces, the fact that tissue-resident populations contain many unique VSGs suggests that blood re-entry may represent a bottleneck for the parasite. An increased rate of switching in tissues could be explained by a higher proportion of dividing slender form parasites in these spaces, as has been observed in the adipose tissue^13^, but we found no correlation at the population level between PAD1 expression and increased VSG diversity. Notably, increased diversity is still observed when the overall parasite load in tissues is lower than the blood (Extended Data Fig 3c). It is therefore exciting to speculate that some aspect of the extravascular environment supplies a molecular or physical stimulus that promotes VSG switching.

While the timing of VSG switching could be the result of an environmental trigger, our data suggest that the higher antigenic diversity in extravascular spaces compared to the blood can also be explained, at least in part, by the dynamics of the immune response to *T. brucei* in each space. In tissue spaces, we observed slower VSG-specific clearance of parasites than in the blood, potentially providing these populations more time to undergo antigenic variation. Additionally, newly switched parasites are still vulnerable to immune clearance by antibodies against their previous VSG for ∼29 hours, so moderate delays in immune clearance could allow more switched parasites to survive^48^. The direct relationship we observed between the timing of parasite clearance and antigenic diversity in AID^-/-^ mice supports this model, with even small delays in clearance showing major effects on parasite VSG diversity in both the tissues and the blood.

Our results show that the production of IgG plays a key role in clearing parasites from tissues, which we propose could be related to the ready diffusion of this molecule within extravascular spaces facilitating parasite clearance. It is important to note, however, that while the difference in timing between the anti-VSG IgM and IgG responses inspired the hypothesis that IgG might be important for parasite clearance in tissues, our study does not prove that this difference in timing explains the delayed clearance in tissue spaces. The local immune response to *T. brucei* is complex, with multiple mechanisms likely to play a role in parasite detection and clearance; our results suggest that IgG is an important player in this response. The increased VSG diversity we observe is certainly multifactorial and could be influenced by parasite factors, such as metabolism, division, motility, and antibody internalization, and host factors, such as extravascular environmental stresses, the local immune response, and vascular permeability. More research will be required to fully understand the complex nature of the *T. brucei* host-pathogen interaction within the extravascular niche.

Altogether, our results outline a model in which *T. brucei* parasites “hide” in extravascular spaces to generate new antigenic variants capable of exiting tissues and aiding in systemic immune evasion. Coupled with other recent studies^9–13,46,49^, this suggests a framework for the progression and pathogenesis of *T. brucei* infections where, instead of being the primary parasite reservoir, the blood may represent a transient population that is regularly re-seeded by extravascular parasites. The vasculature, then, might act as a highway system for movement between the tissue spaces and for eventual transmission back into the tsetse fly.

Interfering with the egress from or establishment within tissue spaces might be a strategy for treating *T. brucei* infections or other infections with pathogens that rely on the distinct features of the extravascular environment. In line with this, one recent study showed that partial inhibition of *T. brucei* tissue invasion using antibodies against P- and E-selectins results in prolonged survival in mice^13^. This also fits with data from another group that found immotile *T. brucei* parasites, likely unable to invade tissues, were no longer infectious^50^. It is possible that without the proper establishment of parasite tissue reservoirs, overall antigenic diversity is lowered, limiting the parasite’s capacity for immune evasion and leading to a decrease in parasite burden. Defining the dynamics and variation of parasites both within and between spaces, as well as the unique host environment within each tissue space, will be central to understanding how *T. brucei* consistently avoids immune clearance and harnessing this mechanism for disease control.

More broadly, these data demonstrate how different environmental and immune pressures within a host can influence pathogen diversification. This production of genetic heterogeneity within an infection is important for many pathogen virulence processes, including establishing and maintaining infection, facilitating immune evasion, generating drug resistance, and adapting to different host environments. The extravascular environment plays a unique role in promoting pathogen evolution, and *T. brucei* serves as a valuable model for understanding this aspect of the host-pathogen interface.

## Methods

### Intravenous mouse infections and sample collection

Female C57Bl/6J (WT, strain# 000664 Jackson Laboratory) or B6.129P2-*Aicda^tm^*^1^(cre)*^Mnz^*/J (AID^-/-^, strain# 007770 Jackson Laboratory)^42^ between 7-10 weeks old were each infected by intravenous tail vein injection with ∼5 pleiomorphic EATRO 1125 AnTat1.1E 90-13 *T. brucei* parasites^24^. Blood parasitemia was counted by tail bleed every 2 days starting on day 4 post-infection (PI) by hemocytometer with a limit of detection of 2.22×10^5^ parasites/mL. Blood (25uL) was collected by a submandibular bleed on days 6, 10, and 14 PI and placed into TRIzol LS. For WT mice, 4 Mice were anesthetized and perfused at days 6, 10, and 14 PI. Infected AID^-/-^ mice were anesthetized and perfused at days 6 (2 mice) and 13 (3 mice) PI. Mice were perfused with 50mL of PBS-Glucose (0.055M D-glucose) with heparin. After perfusion, tissues were dissected and placed immediately into 1mL of RNA Later. The heart, lungs, gonadal fat, subcutaneous fat, brain, and skin (ear) were collected.

For flow cytometry, immunofluorescence experiments, and single-cell sorting experiments, 7-10 week old female C57Bl/6J mice were infected by intravenous tail vein injection with ∼5 AnTat1.1E chimeric triple reporter *T. brucei* parasites which express tdTomato^35^. Blood was collected by a submandibular bleed at designated timepoints. For flow cytometry, mice were anesthetized and perfused on days 6 and 13 P.I. as discussed above and the gonadal fat and lungs were harvested. For immunofluorescence experiments, perfused tissues were collected at day 13 P.I. For single-cell sorting, blood and tissues were only collected on day 14 P.I. All animal studies were approved by the Johns Hopkins Animal Care and Use Committee (protocol # MO22H163).

### VSG-seq sample and library preparation

RNA was isolated from blood samples stored in TRIzol LS (ThermoFisher, 10296010) by phenol/chloroform extraction. Tissue samples were weighed and homogenized in TRIzol, and then RNA was isolated by phenol/chloroform extraction. RNA from each sample was DNase treated using Turbo DNase and cleaned up with Mag-Bind® TotalPure NGS beads (Omega Bio-Tek M1378-00). First-strand cDNA synthesis was performed using SuperScript III Reverse Transcriptase and a primer that binds to the conserved VSG 14-mer in the 3’-UTR (5’-GTGTTAAAATATATC-3’). Products were cleaned up using Mag-Bind® TotalPure NGS beads (Omega Bio-Tek, M1378-01). Next, a VSG-specific PCR with Phusion polymerase (ThermoFisher, F530L) was performed using primers for the spliced leader (5’-ACAGTTTCTGTACTATATTG-3’) and SP6-VSG 14-mer sequences (5’-GATTTAGGTGACACTATAGTGTTAAAATATATC-3’) for 25 cycles. VSG-PCR products were cleaned up using Mag-Bind® TotalPure NGS beads and quantified using the QuBit HS DNA kit (Life Technologies). Finally, sequencing libraries were prepared with the Nextera XT DNA Sample Prep Kit (Illumina) using the manufacturer’s guidelines, and libraries were sequenced with 100bp single-end reads on an Illumina HiSeq 2500.

### Tissue-load and PAD1 QPCRs

First-strand synthesis was performed with SuperScript III Reverse Transcriptase (Thermo Fisher Scientific, 18080051) and random hexamers primers on tissue RNA samples. QPCR was performed in triplicate using SYBR Green qPCR Master Mix (Invitrogen, 4309155). tbZFP3 primers were used to estimate parasite load in tissue samples (FW: 5’-CAGGGGAAACGCAAAACTAA-3’; RV: 5’-TGTCACCCCAACTGCATTCT-3’). CT values were averaged between the triplicates and parasite load per mg of tissue were estimated using a standard curve of values from RNA isolated from known numbers of cultured parasites (Standard curves can be found in Extended Data Fig. 3d).

For PAD1 expression quantification, RNA extraction, first-strand synthesis, and QPCR were performed following the same methods as above. PAD1 expression was quantified by normalizing to tbZFP3 as a control gene (same primers as above) (PAD1 primers; FW: 5’-CAGCGGCGATTATTGCATTGG-3’; RV: 5’-AGGAAGAAGGTTCCTTTGGTC-3’). CT values were averaged between the triplicates and samples were compared using the delta-CT between PAD1 and tbZFP3.

### VSG-seq analysis

Analysis of sequencing results was performed following the method we reported previously^7^, with two changes: no mismatches were allowed for bowtie alignments and each sample was analyzed (assembly, alignment, and quantification) separately. To compare expressed VSG sets between samples, all assembled VSGs were clustered using CD-HIT-EST^51^. VSGs with >98% identity to one another were conservatively treated as one VSG. VSGs were then identified by their Cluster number for further analysis. Samples that had less than 100,000 successfully aligning reads to VSGs were excluded from further analysis. Four samples, 3 brain and 1 heart, were discarded because fewer than 100,000 reads aligned to VSG (Extended Data Fig. 3a). Downstream analysis of expression data and generation of figures was performed in R.

### Analysis of VSG sequence motifs

To identify whether there were tissue-specific VSG sequence motifs, the similarity of N terminal sequences from all assembled VSGs were compared. N terminal sequences were identified using a HMMEr scan against a database curated by Cross et al^2,52^. All N termini were compared in an all vs all blast using default parameters. All VSG pairwise comparisons with an e-value higher than 1E-3 were considered sufficiently similar to one another for further analysis. VSGs that were found in a given tissue were binned into that tissue group, and the distribution of the BLAST bitscores in a given compartment was compared against the total population of similar VSGs.

### Flow Cytometry

Once mice were perfused, tissues were dissected and washed with HBSS (Hanks balanced salt solution, ThermoFisher Scientific 14175095). Tissue samples were minced and placed in DMEM (ThermoFisher Scientific, 11995065) containing either 1 mg/mL collagenase type 1 (ThermoFisher Scientific, 17100017) for adipose fat or 2 mg/mL collagenase type 2 (ThermoFisher Scientific, 17101015) for lung samples. Hearts were dissociated using 2 mg/mL collagenase type 2, 50U/mL DNase I, and 20U/mL Hyaluronidase. These were then incubated in a 37°C water bath for 1 hour and briefly vortexed every 10 minutes. Next, samples were passed through a 70µM filter and centrifuged at 2600 x g for 8 mins at 4 C, and the cell pellet was taken for antibody staining.

Blood samples were collected by submandibular bleed and red blood cells were depleted by magnetic-activated cell sorting (MACS) with anti-Ter-119 MicroBeads (Miltenyi Biotech, 130-049-901) following the manufacturer’s protocol. Cells were pelleted and washed with HMI-9 media.

All samples, both blood and tissues, were stained with Zombie Aqua™ dye at 1:100 in PBS and washed with PBS following the manufacturer’s protocol (BioLegend, 423101). Samples were then stained for 10 minutes at 4°C with a rabbit anti-AnTat1.1 polyclonal antibody diluted 1:15,000 in HMI-9 media and washed once with HMI-9 (antibody courtesy of Jay Bangs). Then, secondary antibody staining was performed while shaking for 10 minutes at 4°C with Anti-Rabbit IgG (H+L), F(ab’)2 Fragment conjugated to Alexa Fluor® 488 fluorescent dye (Cell Signaling Technology, 4412S). Finally, samples were washed with cold PBS and resuspended in PBS for flow cytometry analysis. Samples were run on a Beckton Dickenson A3 Symphony flow cytometer and analysis was performed using FlowJo (version 10.6.1) (see Extended Data Fig. 7a for gating strategy).

### Immunofluorescence

Mice infected with AnTat1.1E chimeric triple reporter *T. brucei* parasites that express tdTomato were sacrificed and perfused as previously described at days 6 and 13 PI. Lung, heart, and gonadal fat were collected and fixed in 4% paraformaldehyde in PBS for 12 hours at 4 C. Post-fixation, tissues were frozen, embedded in O.C.T. Compound (Tissue-Tek), and cut by cryostat microtome into 10μm sections.

The following antibodies were applied to sections: rat anti-mouse CD31 (PECAM-1) (SCBT) with goat anti-rat Fluor 488 (CST). Coverslips were mounted using ProLong Gold (Life technologies). Tissues were imaged with 4x, 10x, and 20x objectives using a Nikon Eclipse 90i fluorescence microscope (Nikon) and X-Cite 120 fluorescent lamp (Excelitas) with an ORCA-ER digital CCD camera (Hammamatsu) and ImageJ v1.53 image analysis software. Image collection and analysis followed published guidelines for rigor and reproducibility^53^.

### Serum antibody ELISA quantification

Blood (25uL) was collected by submandibular bleed on days 0, 6, 10, and 14 PI from mice infected with ∼5 pleiomorphic EATRO 1125 AnTat1.1E 90-13 *T. brucei* parasites. Serum was isolated using serum separator tubes (BD Microtainer SST tubes, 365967). IgM and IgG were quantified by ELISA using ThermoFisher IgM and IgG kits following manufacturer protocols (IgG cat# 88-50400-88, IgM cat# 88-50470-88).

### Serum sample flow cytometry on *T. brucei*

Blood was collected from 2 mice infected with AnTat1.1E chimeric triple reporter *T. brucei* parasites, which initially express the VSG AnTat1.1, by cheek bleed. Serum was isolated by spinning blood samples at 10,000xg for 5 mins and pipetting off the top serum layer. EATRO 1125 AnTat1.1E 90-13 *T. brucei* parasites^24^ expressing the VSG AnTat1.1 and Monomorphic Single Marker Lister427 VSG221 TetR T7RNAP bloodstream form (NR42011; LOT: 61775530)^54^, which express VSG-2, were used for flow cytometry. 10^6^ Parasites were stained in duplicate while shaking for 10 minutes at 4°C using mouse serum diluted 1:100 in PBS. As a positive control, a rabbit anti-AnTat1.1 polyclonal antibody diluted 1:15,000 in PBS was used following the same staining procedure (antibody courtesy of Jay Bangs). Then, secondary antibody staining was performed while shaking for 10 minutes at 4°C with Anti-mouse IgG (H+L), F(ab’)2 Fragment conjugated to Alexa Fluor® 647 fluorescent dye (Cell Signalling Technology, 4410S) or Anti-Rabbit IgG (H+L), F(ab’)2 Fragment conjugated to Alexa Fluor® 647 fluorescent dye (Cell Signaling Technology, 4414S). Finally, samples were washed with cold PBS and resuspended in PBS for flow cytometry analysis. Samples were run on an Attune Nxt flow cytometer (Invitrogen) and analysis was performed using FlowJo (version 10.6.1).

### Single-cell sorting and RNA-seq library preparation

Blood and tissue samples (heart, gonadal fat, and lung) from two mice were collected after perfusion as described above. For sorting parasites, the same tissue dissociation and flow cytometry was performed as described above, with the exception of live/dead staining, which was performed using propidium iodide instead of Zombie Aqua™. Samples were kept on ice as much as possible through this process. Single live, tdTomato^+^ *T. brucei* cells were sorted into chilled 384-well plates for SL-Smart-seq3xpress library preparation containing lysis buffer and an RNA spike-in control using a Beckman Coulter MoFlo™ XDP cell sorter (see Extended Data Fig. 7b for gating strategy). For each blood and tissue sample a single plate of parasites were sorted for a total of 370 cells per sample.

The SL-Smart-seq3xpress library preparation approach was followed as described in McWilliam et al^33^. Briefly, single cells were lysed by incubation at 72°C for 10 minutes in 0.3 µl of lysis buffer. Reverse transcription was done by adding 0.1µl of reverse transcription mix to each well and incubation at 42°C for 90 minutes, followed by ten cycles of 50°C for 2 minutes and 42°C for 2 minutes, with a final incubation at 85°C for 5 minutes. Preamplification was done by adding 0.6µl of a mix containing primers annealing to the Spliced-Leader sequence and to a conserved sequenced added by the oligodT primer during retrotranscription, with the following cycling conditions: 95°C for 1 minute, 16 cycles of : 98°C for 10 seconds, 65°C for 30 seconds, 68°C for 4 minutes; and final 72°C for 10 minutes. Following, the amplified cDNA was diluted by adding 9µl of water per well. Next, 1 µl of each well was transferred to a new plate, 1µl of tagmentation mix was added and the plate incubated at 55°C for 10 minutes for tagmentation. The reaction was stopped by adding 0.5µl of 0.2% SDS to each well and incubating for 5 minutes. The final index PCR was done by adding 1µl of specific index primer combinations to each well and 1.5µl of PCR mix. The following cycling conditions were used: 72° for 3 minutes, 95°C for 30 seconds, 14 cycles of: 95°C for 10 seconds, 55°C for 30 seconds, 72°C for 1 minute; followed by 72°C for minutes. Single-cell libraries were then pooled and purified using AMPure XP beads at a ratio of 1:0.7. Libraries were run on a 4% non-denaturing PAGE gel and purified according to standard polyacrylamide gel purification protocols. Purified libraries from multiple plates were pooled and sequenced on a NextSeq 1000 sequencing platform to produce paired-end reads of 101nt (cDNA) and 19nt (TAG+UMI read), and 8nt for the index reads.

### Single-cell RNA-seq primary processing and VSG de novo assembly

The primary processing of the sequencing data was as described in McWilliam et al^33^. Briefly, the two reads containing the indexes (8nt each) and the one containing the TAG+UMI (19nt) were concatenated into a 35nt read. Artifact reads containing the TAG sequence (or its reverse complement) in the cDNA reads were filtered out with Cutadapt.

For analysis of derepression by *de novo* assembly of VSGs, the filtered reads were sorted into individual files for each cell, and these read files were run through our VSG-Seq analysis pipeline individually using the same parameters as described above for bulk VSG-Seq. Using this approach, VSG open reading frames (ORFS) were assembled and quantified for each cell individually. VSG ORFs were then clustered among all single cells (VSG clusters were not related to previous VSG clusters from bulk VSG-seq and cannot be compared to the previous analysis based on cluster names).

For analysis of derepression by alignment to the genome, filtered reads were mapped with STAR (version 2.7.10a) to a hybrid fasta file combining the *T. brucei* EATRO 1125 strain genome assembly (version 67, downloaded from TriTrypDB^55^) and the set of 10 sequences used as RNA spike-in. The count matrix obtained was then corrected with the index hopping filtering pipeline “scSwitchFilter” (https://github.com/colomemaria/scSwitchFilter). Only cells with at least 500 genes detected, 1000 gene UMI transcript counts, 30 spike-in UMI counts, and 10 VSG UMI counts were used for downstream analyses. A total of 1216 total cells fit these criteria out of 2960 total cells sequenced. For each tissue and blood sample a single plate (370 cells) was sequenced. For assessment of potential derepression (Extended Data Figure 5b,e), VSGs with >1 UMI count were considered expressed in a cell. Cells were considered to be monogenically expressing a VSG (Extended Data Figure 5e) if the VSG represented ≥80% of VSG UMI counts.

For evaluating read coverage of VSGs and VSG clusters, Bowtie indexes were created for each reference sequence then reads from a single cell were aligned using Bowtie1. Read coverage was calculated using deepTools^56^ to convert BAM alignment files to bigWig coverage tracks. Coverage was then visualized using the ggcoverage package in R (https://github.com/showteeth/ggcoverage).

### Tsetse fly husbandry and fly bite-inoculated mouse infections

*Glossina morsitans morsitans* were maintained in the Yale School of Public Health insectary at 25°C with 65-70% relative humidity under a 12h:12h light:dark photoperiod. All flies received defibrinated sheep blood (Lampire Biologicals) every 48 hours through an artificial membrane feeding system^57^. Newly eclosed adult female flies were administered *per os* an initial blood meal containing 1×10^6^/ml of *Trypanosoma brucei brucei* (strain RUMP 503; previously expanded in rats) and cysteine (10µM; to increase the infection prevalence; ^58^). After this single parasite challenge, flies were maintained on normal blood every other day.

Thirty-five days post-challenge (the time it takes *T. b. brucei* to complete their developmental cycle within the tsetse fly and become infectious to a new vertebrate host), six to eight-week-old female C57Bl/6J mice were exposed to the bite of individual, trypanosome challenged flies 72 hrs after the flies had taken their last blood meal. Following the consumption of mouse blood, individual flies were microscopically dissected to confirm that their salivary glands were infected with vertebrate-infectious metacyclic stage *T. b. brucei* (if not, another fly was allowed to feed on the mouse until it was confirmed that an infectious fly had taken a blood meal). Five mice were infected using this method. All experiments using mice were performed in strict accordance with the Yale University Institutional Animal Care and Use Committee policies (Protocol 2014–07266 renewed on March 2023).

Once mice were infected, blood parasitemia was counted by tail bleed every 2 days starting on day 4 post-infection (PI) by hemocytometer with a limit of detection of 2.22×10^5^ parasites/mL. Blood (25uL) was collected by a submandibular bleed on days 6, 10, and 14 PI and placed into TRIzol LS. Five Mice were anesthetized and perfused at day 14 PI. Mice were perfused with 50mL of PBS-Glucose (0.055M D-glucose) with heparin. After perfusion, tissues were dissected and placed immediately into 1mL of RNA Later. The heart, lungs, gonadal fat, subcutaneous fat, brain, and skin (ear) were collected. Sequencing libraries were prepared and analyzed following the methodology described above.

To quantify the mVSG repertoire of RUMP 503, we also collected a pool of tsetse saliva containing RUMP 503 *T. brucei* parasites. This sample was stored in TRIzol LS, RNA was extracted, and the sample was prepared for sequencing as described above. The VSG-Seq pipeline was used to quantify mVSG expression in the sample.

### Statistics and figures

Normality was tested for all Students t-tests and Dunnet’s tests analyses and can be found in the code on the repository at https://github.com/mugnierlab/Beaver2022. Nearly all VSG diversity measurements were found to be normally distributed, except for some samples with low VSG counts or sample numbers. We thus assumed normality for all VSG diversity measurements. All reported P-values have been corrected for multiple comparisons using the Benjamini–Hochberg procedure. For all figures with boxplots, the box represents the first (25%) and third (75%) quartiles with a line at the median. Extending lines represent the maximum and minimum values not including outliers that are further than 1.5 times the interquartile range from either end of the box.

## Data availability

Code and data for generating the analysis and figures in this paper are available at https://github.com/mugnierlab/Beaver2022. Raw sequencing data are available in National Center for Biotechnology Information (NCBI) Sequence Read Archive under accession number PRJNA858046.

## Acknowledgments

We would like to thank Jay Bangs for supplying us with the anti-AnTat1.1 antibody and Brice Rotureau for providing us with the chimeric triple reporter cell line. We thank Tricia Nilles and Worod Allak at the Becton Dickinson Immunology and Flow Cytometry core at Johns Hopkins Bloomberg School of Public Health for training, support, and technical assistance using the BD FACS Symphony A3. We would like to also thank Hao Zhang and Joseph Margolick from the Flow Cytometry Cell Sorting Core Facility at Bloomberg School of Public Health, Johns Hopkins University for doing FACS sorting. The facility was supported by CFAR: 5P30AI094189-04 (Chaisson), 1S10OD016315-01 and 1S10RR13777001. Data analysis was carried out at the Advanced Research Computing at Hopkins (ARCH) core facility (rockfish.jhu.edu), which is supported by the National Science Foundation (NSF) grant number OAC 1920103. AKB was supported by NIH T32AI007417. NPC was supported by NIH T32 OD011089. FR-F is supported by NIH NIGMS 1K99GM132557-01 and is an Investigator in the Chan Zuckerberg Biohub. MRM, AKB, JES, NPC, JMCH, BB, and BZ were supported by Office of the Director, NIH (DP5OD023065). In addition, this study was funded by the German Research Foundation [SI 1610 / 2-2] and an ERC Consolidator Grant (SwitchDecoding 101044320) awarded to TNS. ZK was supported by a MSCA ITN Cell2Cell fellowship.

## Author contributions

Conceptualization: AKB, MRM, LMF, FRF, TNS, ROC, SA, BLW

Methodology: AKB, ROC, ZK, BLW, EOA, GMS, NPC, GYB, JMCH, JES, BZ, BB

Investigation: AKB, ROC, ZK, BLW, NPC, GYB, JH, JES

Visualization: AKB, NPC, JMCH

Funding acquisition: MRM, LMF, TNS, SA

Project administration: MRM

Supervision: MRM

Writing – original draft: AKB, MRM

Writing – review & editing: AKB, JES, GYB, BZ, MRM, FRF, LMF, TNS, ROC

## Materials and correspondence

Correspondence and requests for materials should be addressed to Monica Mugnier.

## Extended data

**Extended Data Fig. 1.**
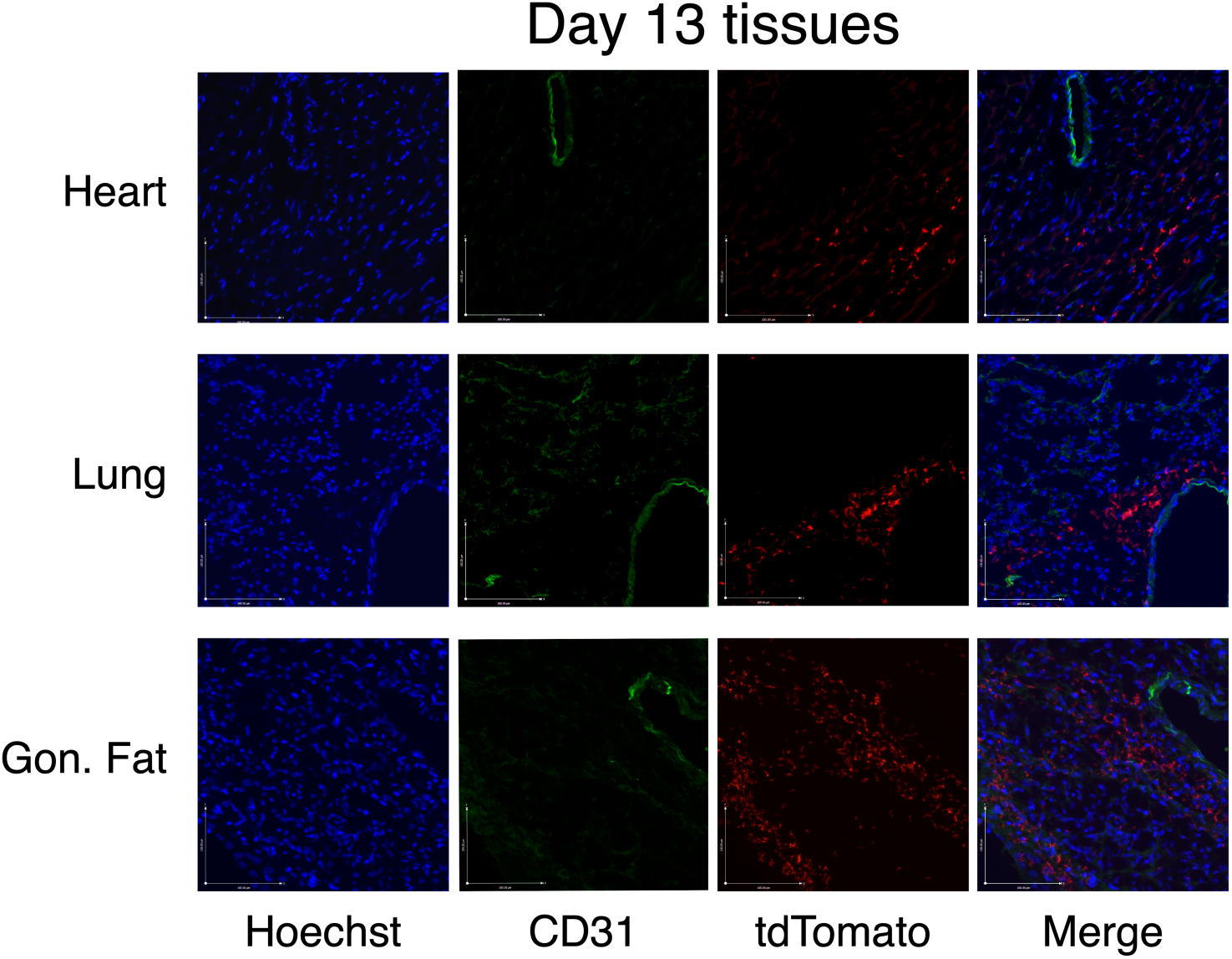
Immunofluorescence images of tdTomato expressing parasites (red) from cross sections of perfused tissues stained with hoechst (blue) and anti-CD31 antibody (green). TdTomato-positive parasites localized separately from CD31 lined spaces, showing that parasites are extravascular.

**Extended Data Fig. 2.**
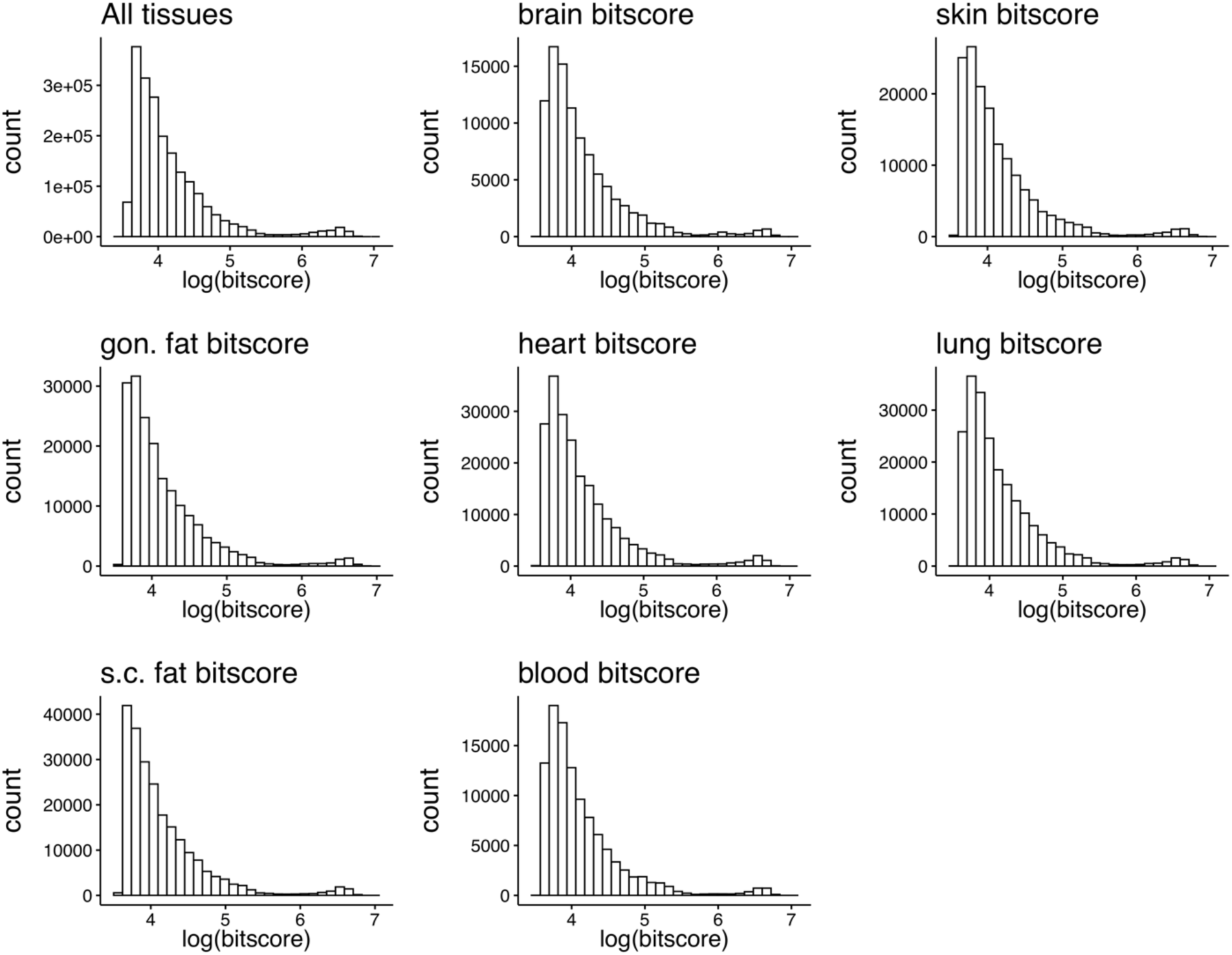
Distribution of bitscores across tissue compartments. The similarity of VSGs detected in each tissue as measured by bitscore, a sequence similarity metric normalized to the database size, allows for comparing tissue compartments with different numbers of total VSGs expressed in each tissue. No statistical significance was found.

**Extended Data Fig. 3.**
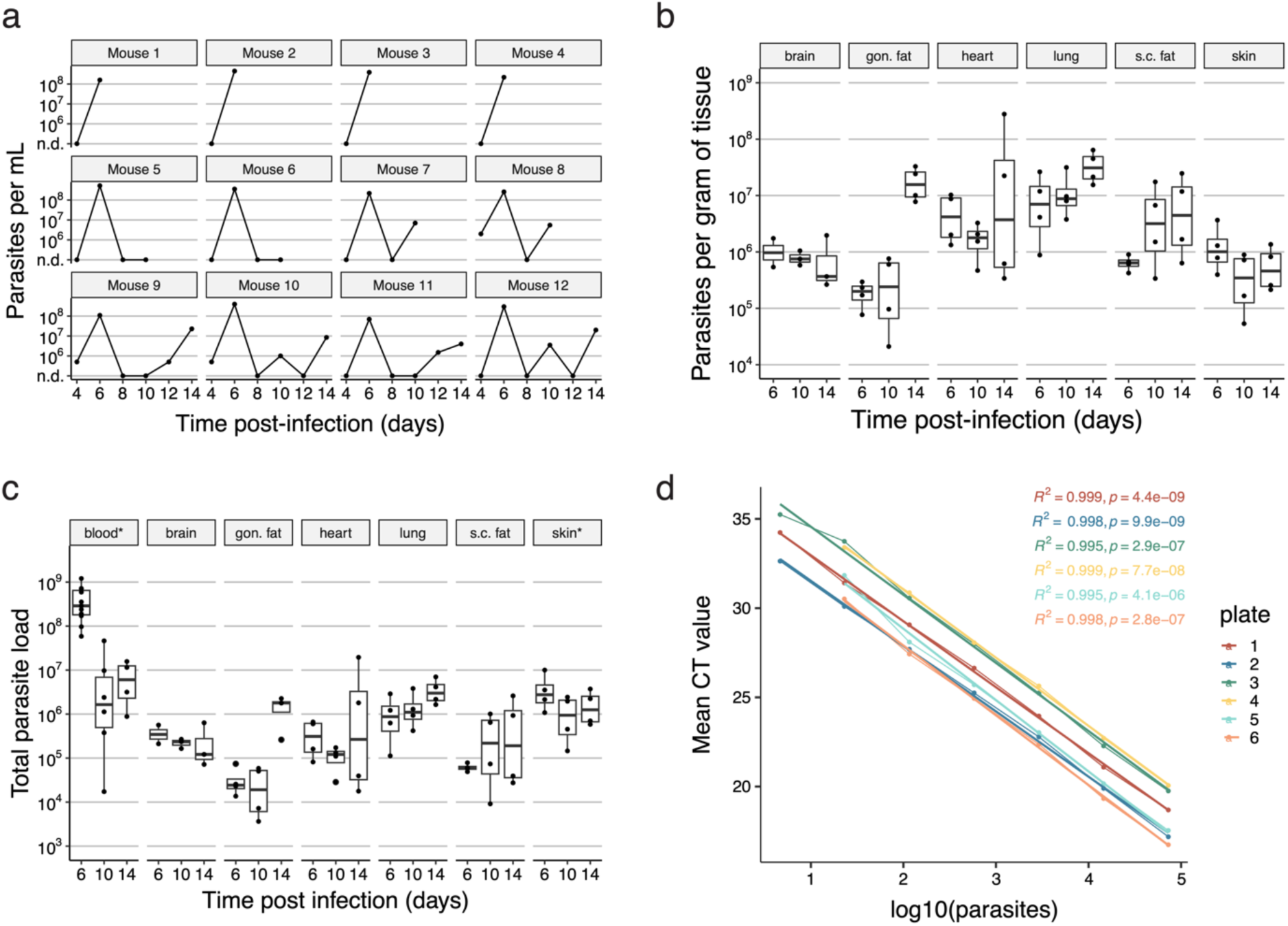
**(a**) Parasitemia of 12 mice infected with AnTat1.1E *T. brucei* counted from tail blood by hemocytometer (“n.d.” = not detectable, limit of detection of 2.22×10^5^ parasites/mL). (**b**) Estimated parasite load per gram of tissue using QPCR. tbZFP3 was used as the control and RNA from known quantities of parasites was used to make standard curves. (**c**) The approximate total number of parasites represented in each organ. This was calculated using the estimated number of parasites from QPCR and the recorded organ mass. For the blood and skin, it was assumed that each mouse had 1.5mL of blood and 2.73 grams of skin^59^ to estimate the total load within these organs. (**d**) The qPCR standard curves used for each plate of samples. These were used to estimate the number of parasites represented in each of our tissue samples based on RNA from known parasite concentrations (cultured parasites counted using a hemocytometer).

**Extended Data Fig. 4.**
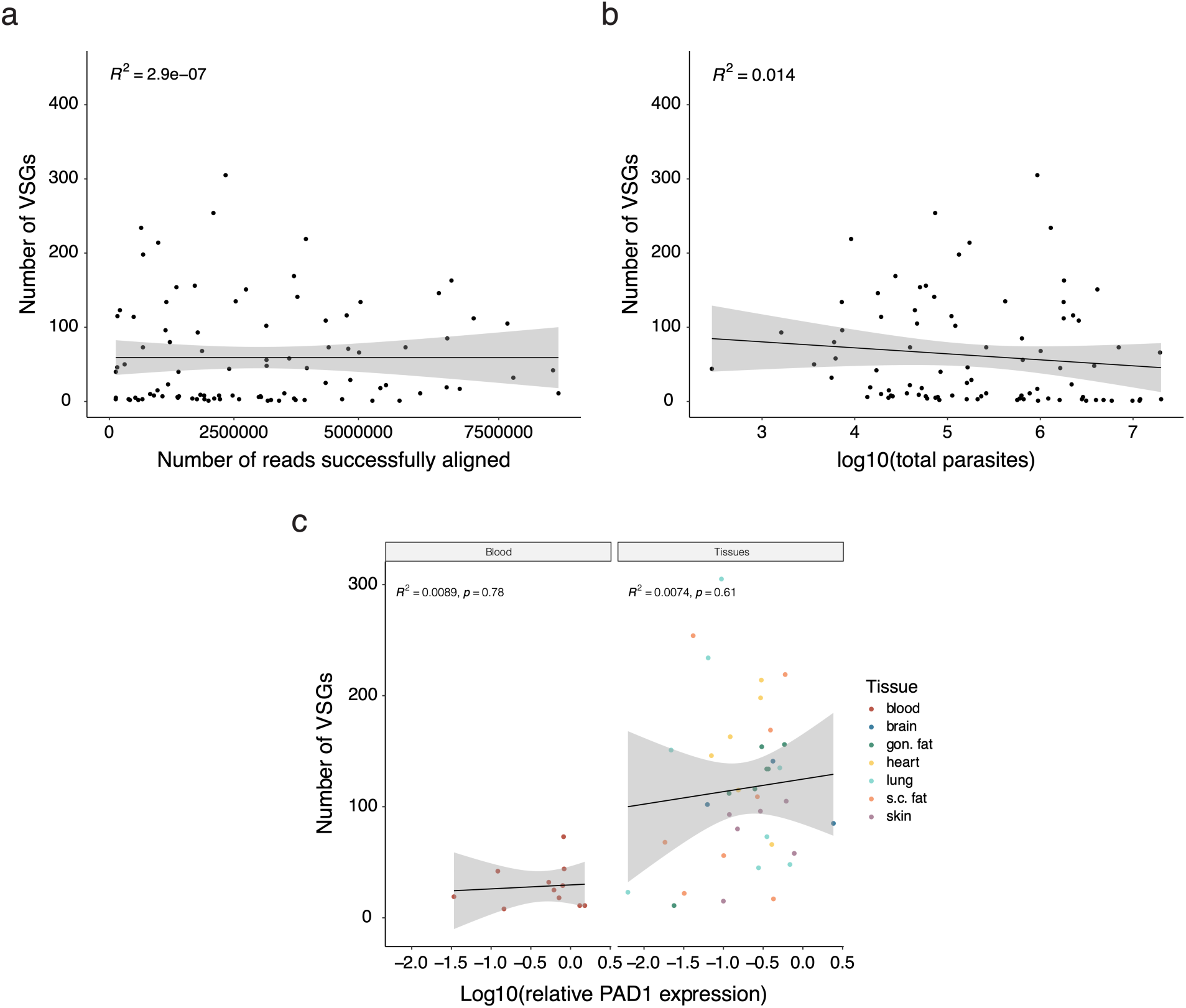
(**a**) A comparison of the number of reads successfully aligned in a sample and the number of VSGs observed. (**b**) A comparison of the total number of parasites and the number of VSGs found in each sample. (**c**) The correlation between PAD1 expression relative to the housekeeping gene tbZFP3 and the number of VSGs expressed at the population level for each sample.

**Extended Data Fig. 5.**
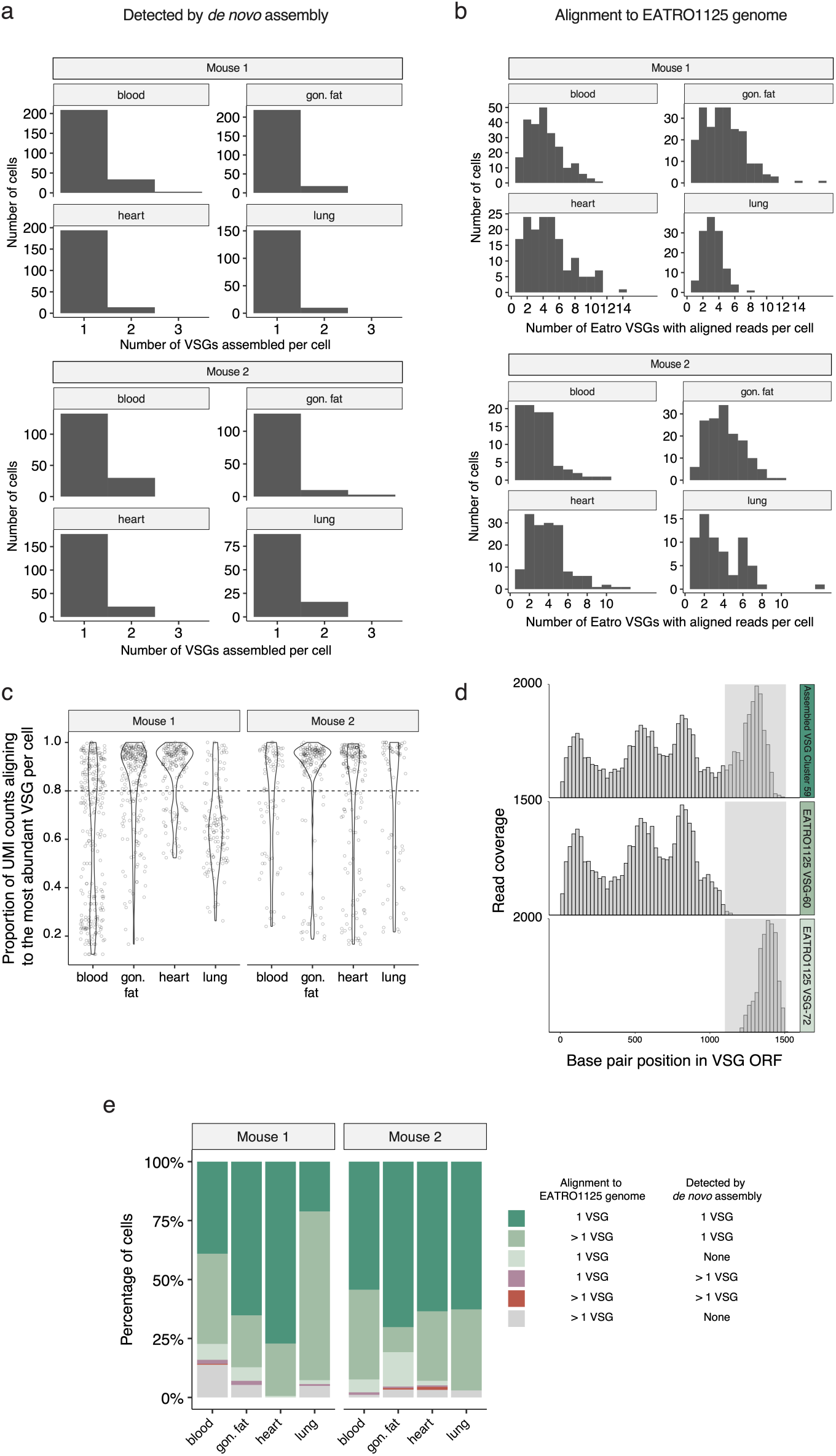
(**a**) The number of VSG open reading frames (ORFs) detected per cell by *de novo* assembly with the VSG-Seq pipeline. In this figure, all cells sequenced were evaluated for VSG ORF assembly without any filtering (1377 total cells had a *de novo* assembled VSG). (**b**) The number of EATRO1125 VSGs detected per single cell. Cells were evaluated for VSG expression only if they had at least 500 genes detected, 1000 gene UMI counts, 30 spike-in UMI counts, and 10 VSG UMI counts (1216 cells fit these criteria out of 2960 total sequenced cells). Only VSGs with >1 UMI count were considered for quantification of VSG expression. (**c**) Alignment to the EATRO1125 genome assembly was used to quantify the fraction of VSG UMI counts coming from the most abundant VSG gene within each cell. The dashed line represents the 0.8 fraction of total VSG UMI counts (80%) threshold set to define monogenic expression with one dominant VSG in a cell. Only cells that had at least 500 genes detected, 1000 gene UMI transcript counts, 30 spike-in UMI counts, and 10 VSG UMI counts were evaluated for this analysis (1216 total cells out of 2960 total cells sequenced). (**d**) Representative histograms of coverage for reads from one cell. Read coverage is shown for the *de novo* assembled VSG cluster 59 and two genomic VSG ORFs, VSG-60 and VSG-72, which represent a common scenario that creates ambiguous VSG expression if reads are only aligned to the EATRO1125 genome. Many cells in the Mouse 1 lung sample expressed this VSG and exhibited this mapping problem. (**e**) The VSG expression classification for each cell using both methods (alignment to the EATRO1125 genome and *de novo* assembly). In both alignment and assembly, if a VSG represented 80% or more of the VSG UMI counts in a cell (for genomic mapping) or 80% of the population (for VSG-seq analysis), that cell was considered to be expressing only the dominant VSG. Only cells that had at least 500 genes detected, 1000 gene UMI transcript counts, 30 spike-in UMI counts, and 10 VSG UMI counts were evaluated for this analysis (1216 cells fit these criteria out of 2960 total cells sequenced). Only VSGs with >1 UMI count were considered for quantification by alignment to the EATRO1125 genome.

**Extended Data Fig. 6.**
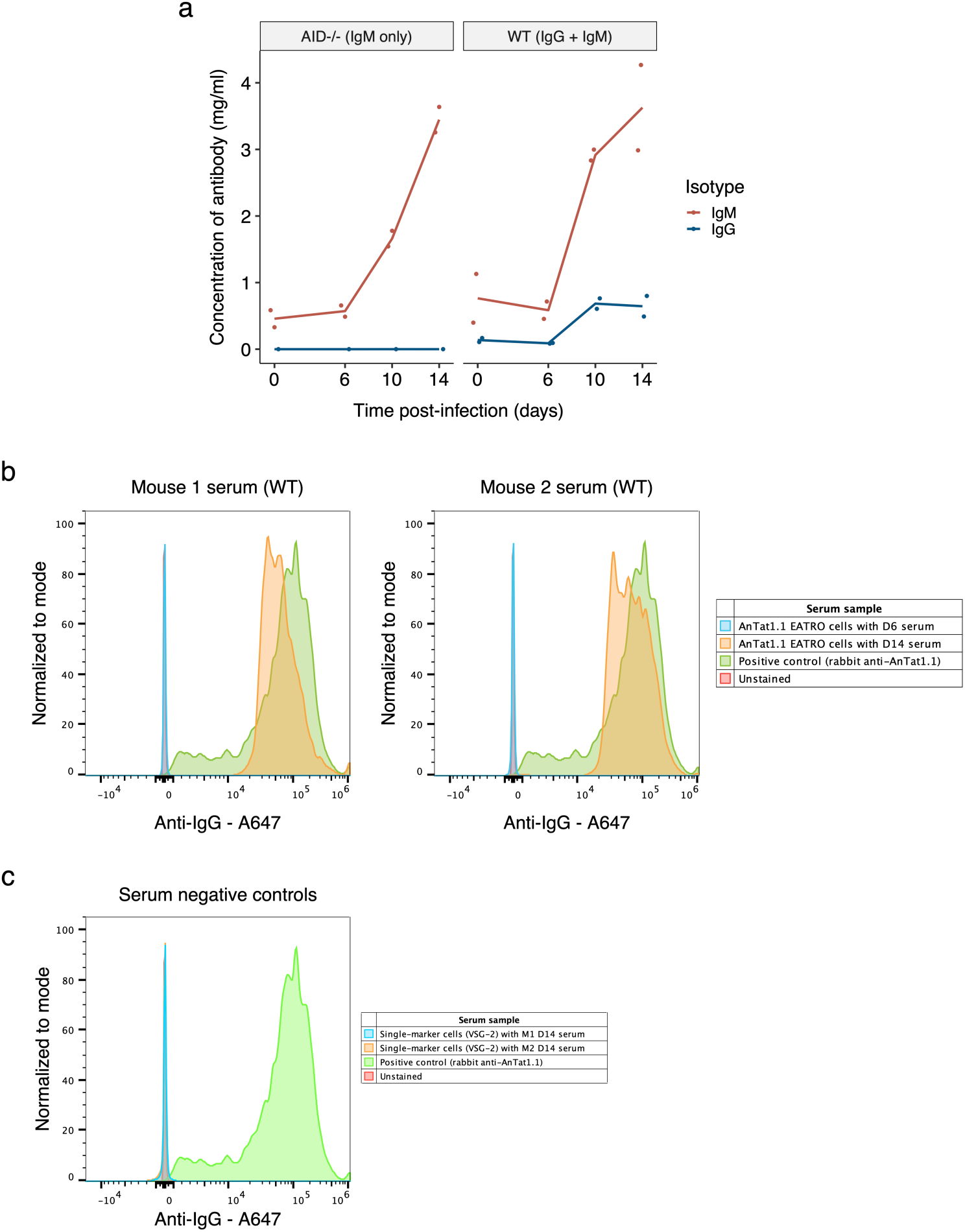
**(a**) Quantification of serum IgM and IgG concentrations in infected AID^-/-^ and wildtype (WT) mice by ELISA (n = 2). (**b**) AnTat1.1 EATRO cells (expressing VSG AnTat1.1) stained with serum, followed by an anti-IgG secondary antibody that cross-reacts with IgM and other isotypes, from day 6 and day 14 of two mice infected with AnTat1.1 Triple-marker cells. A polyclonal rabbit anti-AnTat1.1 antibody on cells known to be expressing AnTat1.1 was used as a positive control. (**c**) Single-marker cells (expressing VSG-2) stained with day 14 serum from two mice infected with AnTat1.1 Triple-marker cells. These were used as negative controls to show that serum from infected mice specifically binds AnTat1.1 and no other VSGs.

**Extended Data Fig. 7.**
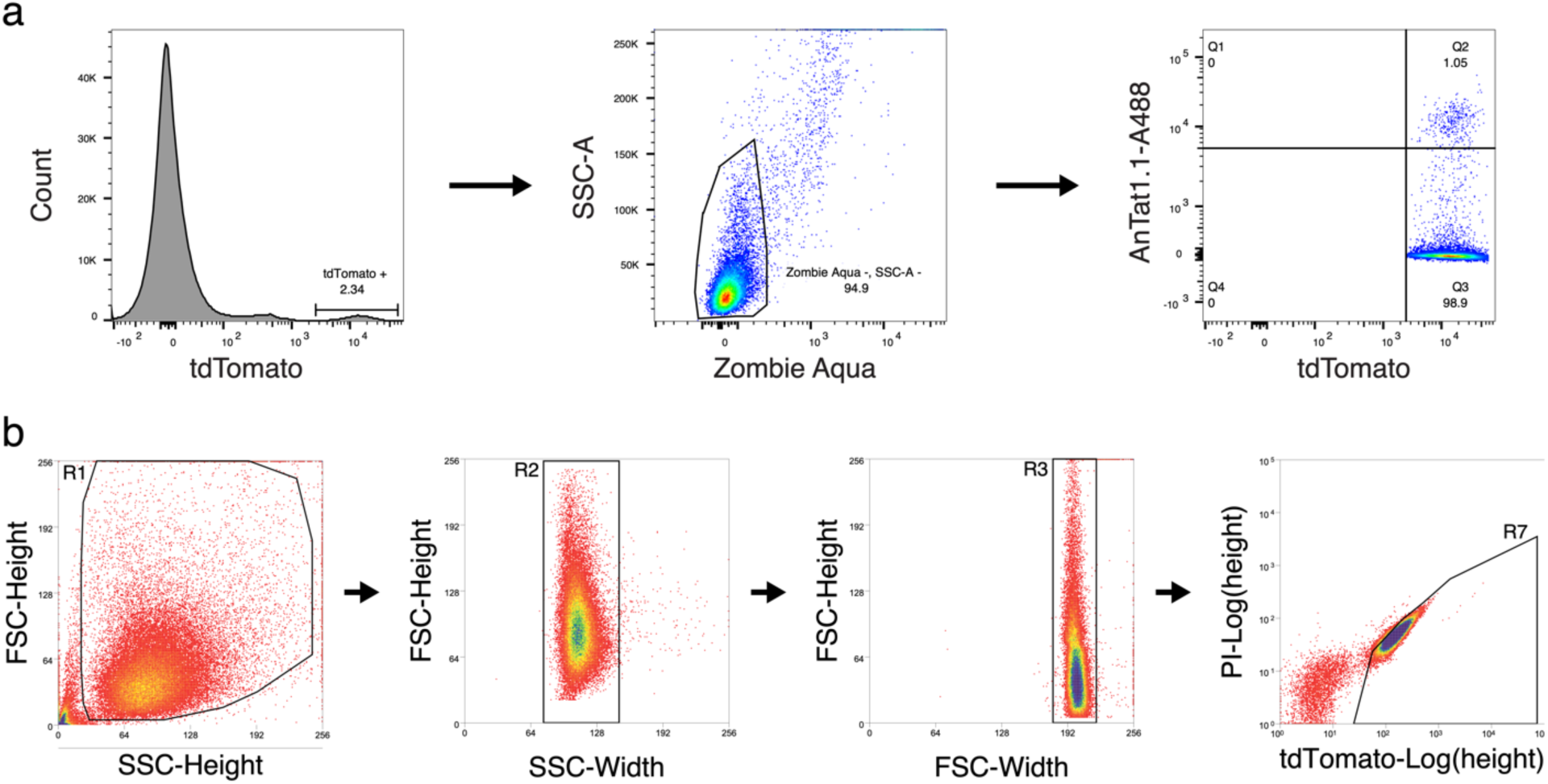
(**a**) Example gating strategy for flow cytometry experiments from Figure 3. Samples were first gated for tdTomato-positive cells, which represent *T. brucei* parasites expressing tdTomato in their cytoplasm. Then live cells were gated based on live/dead Zombie Aqua^TM^ staining. Finally, quadrants were placed around AnTat1.1-A488 positive and negative cells. (**b**) Gating strategy for single-cell sorting into 384-well plates for the SL-Smart-seq3xpress platform. This example is from the blood of mouse 2. Singlet parasites were selected and then tdTomato positive, PI (propidium iodide) negative *T. brucei* cells were sorted into single cells.

**Extended Data Table 1.** Flow cytometry counts of tdTomato-positive parasites from infections with triple-marker *T. brucei* parasites. Parasites were collected from the blood and dissociated tissues of 10 mice on either day 6 or 13 post-infection and stained with an anti-AnTat1.1 polyclonal antibody. Here we report the raw number of live tdTomato-positive parasites that were Antat1.1 positive and negative in each sample. These data were used to generate Fig. 3b.

| *Biological experiment* | *Mouse* | *Tissue* | *Day* | *Count of AnTat1.1 + cells* | *Count of AnTat1.1 - cells* | *Total number of cells* | *Percent of cells AnTat1.1+* |
| --- | --- | --- | --- | --- | --- | --- | --- |
| 1 | 1 | blood | 6 | 20394 | 486 | 20880 | 97.67% |
| 1 | 1 | gon. fat | 6 | 411 | 32 | 443 | 92.78% |
| 1 | 1 | lung | 6 | 817 | 49 | 866 | 94.34% |
| 1 | 2 | blood | 6 | 42969 | 1458 | 44427 | 96.72% |
| 1 | 2 | blood | 13 | 1 | 19922 | 19923 | 0.01% |
| 1 | 2 | gon. fat | 13 | 0 | 250 | 250 | 0.00% |
| 1 | 2 | lung | 13 | 573 | 4460 | 5033 | 11.38% |
| 2 | 3 | blood | 6 | 19737 | 270 | 20007 | 98.65% |
| 2 | 3 | gon. fat | 6 | 220 | 4 | 224 | 98.21% |
| 2 | 3 | lung | 6 | 1464 | 55 | 1519 | 96.38% |
| 2 | 4 | blood | 6 | 39373 | 592 | 39965 | 98.52% |
| 2 | 4 | gon. fat | 6 | 379 | 10 | 389 | 97.43% |
| 2 | 4 | lung | 6 | 676 | 30 | 706 | 95.75% |
| 2 | 5 | blood | 6 | 49406 | 600 | 50006 | 98.80% |
| 2 | 5 | blood | 13 | 1 | 19885 | 19886 | 0.005% |
| 2 | 5 | gon. fat | 13 | 31 | 4982 | 5013 | 0.62% |
| 2 | 5 | lung | 13 | 318 | 29889 | 30207 | 1.05% |
| 2 | 6 | blood | 6 | 10317 | 31 | 10348 | 99.70% |
| 2 | 6 | blood | 13 | 0 | 1001 | 1001 | 0.00% |
| 2 | 6 | gon. fat | 13 | 3 | 781 | 784 | 0.38% |
| 2 | 6 | lung | 13 | 265 | 29733 | 29998 | 0.88% |
| 3 | 7 | blood | 6 | 65135 | 249 | 65384 | 99.62% |
| 3 | 7 | gon. fat | 6 | 69 | 3 | 72 | 95.83% |
| 3 | 7 | lung | 6 | 275 | 182 | 457 | 60.18% |
| 3 | 8 | blood | 6 | 49239 | 632 | 49871 | 98.73% |
| 3 | 8 | gon. fat | 6 | 71 | 0 | 71 | 100.00% |
| 3 | 8 | lung | 6 | 770 | 137 | 907 | 84.90% |
| 3 | 9 | blood | 6 | 49758 | 147 | 49905 | 99.71% |
| 3 | 9 | blood | 13 | 3 | 5042 | 5045 | 0.06% |
| 3 | 9 | gon. fat | 13 | 0 | 1537 | 1537 | 0.00% |
| 3 | 9 | lung | 13 | 403 | 18944 | 19347 | 2.08% |
| 3 | 10 | blood | 6 | 49286 | 642 | 49928 | 98.71% |
| 3 | 10 | blood | 13 | 14 | 20204 | 20218 | 0.07% |
| 3 | 10 | gon. fat | 13 | 33 | 7034 | 7067 | 0.47% |
| 3 | 10 | lung | 13 | 187 | 38096 | 38283 | 0.49% |

**Extended Data Table 2.** Summary of single cell sequencing results. *T. brucei* single cells were sorted from blood and tissues on day 14 post-infection from 2 mice and sequenced using the SL-Smart-seq3xpress platform. For *de novo* assembly of VSGs, all cells were considered without filters, but for all other columns only cells that had at least 500 genes detected, 1000 gene UMI counts, 30 spike-in UMI counts, and 10 VSG UMI counts were considered. Additionally, only VSGs that had >1 UMI counts when aligning to the EATRO1125 genome were considered to be detected.

| Number of single cells |  |  |  |  |  |  |  |  |  |  |  |
| --- | --- | --- | --- | --- | --- | --- | --- | --- | --- | --- | --- |
| Mouse | Tissue | Total cells sequenced | Cells passing QC cutoffs | Average successfully aligning UMI counts per cell | Cells with a de novo assembled VSG ORF | 1 VSG by alignment and 1 VSG by de novo assembly | >1 VSG by alignment and 1 VSG by de novo assembly | 1 VSG by alignment and no VSG by de novo assembly | 1 VSG by alignment and >1 VSG by de novo assembly | >1 VSG by alignment and >1 VSG by de novo assembly | >1 VSG by alignment and no VSG by de novo assembly |
| Mouse 1 | blood | 370 | 238 | 5736 | 227 | 93 | 91 | 16 | 4 | 1 | 33 |
|  | gon. fat | 370 | 227 | 4275 | 228 | 148 | 50 | 13 | 4 | 0 | 12 |
|  | heart | 370 | 162 | 5375 | 201 | 125 | 36 | 1 | 0 | 0 | 0 |
|  | lung | 370 | 123 | 2952 | 156 | 26 | 88 | 2 | 1 | 0 | 6 |
| Mouse 2 | blood | 370 | 92 | 3572 | 148 | 50 | 35 | 5 | 1 | 0 | 1 |
|  | gon. fat | 370 | 151 | 3997 | 133 | 106 | 16 | 22 | 1 | 1 | 5 |
|  | heart | 370 | 156 | 4699 | 188 | 99 | 46 | 3 | 1 | 2 | 5 |
|  | lung | 370 | 67 | 5097 | 96 | 42 | 23 | 0 | 0 | 0 | 2 |
| **Totals** |  | 2960 | 1216 | 4462 | 1377 | 689 | 385 | 62 | 12 | 4 | 64 |

## Supplementary information

**Supplementary Data 1:** A summary of VSG expression in each cell by both genome alignment and *de novo* assembly, including categorization for Extended Data Figure 5e.

**Supplementary Data 2**: The EATRO1125 VSG read alignment in each cell that meets QC cutoffs (at least 500 genes detected, 1000 gene UMI counts, 30 spike-in UMI counts, and 10 VSG UMI counts) and has >1 VSG UMI count. Raw alignment count tables and VSG count tables that include unfiltered cells can also be found at https://github.com/mugnierlab/Beaver2022.

**Supplementary Data 3**: The results from *de novo* assembly of VSGs in each cell. The Trinity name for each ORF and the top VSG blast hit for each assembled VSG ORF is reported, including the proportion of VSG expression accounted for by that VSG. For each cell, a dominant VSG is identified if it represented over 80% of the VSG expressed in the cell and “NA” denotes that there is not a dominant VSG in that cell. VSG assembly was attempted in all cells without any previous QC filtering. Here, all VSG assemblies are reported without further filtering and the sequences can be found at https://github.com/mugnierlab/Beaver2022.

